# The focal adhesion protein talin is a mechanically-gated A-kinase anchoring protein (AKAP)

**DOI:** 10.1101/2023.08.20.554038

**Authors:** Mingu Kang, Yasumi Otani, Yanyu Guo, Jie Yan, Benjamin T. Goult, Alan K. Howe

## Abstract

The cAMP-dependent protein kinase (Protein Kinase A; PKA) is a ubiquitous, promiscuous kinase whose activity is focused and specified through subcellular localization mediated by A-kinase anchoring proteins (AKAPs). PKA has complex roles as both an effector and a regulator of integrin-mediated cell adhesion to the extracellular matrix (ECM). Recent observations demonstrate that PKA is an active component of focal adhesions (FA), intracellular complexes coupling ECM-bound integrins to the actin cytoskeleton, suggesting the existence of one or more FA AKAPs. Using a combination of a promiscuous biotin ligase fused to PKA type-IIα regulatory (RIIα) subunits and subcellular fractionation, we identify the archetypal FA protein talin1 as an AKAP. Talin is a large, mechanosensitive scaffold that directly links integrins to actin filaments and promotes FA assembly by recruiting additional components in a force-dependent manner. The rod region of talin1 consists of 62 α-helices bundled into 13 rod domains, R1-R13. Direct binding assays and nuclear magnetic resonance spectroscopy identify helix41 in the R9 subdomain of talin as the PKA binding site. PKA binding to helix41 requires unfolding of the R9 domain, which requires the linker region between R9 and R10. Finally, single-molecule experiments with talin1 and PKA, and experiments in cells manipulated to alter actomyosin contractility demonstrate that the PKA-talin interaction is regulated by mechanical force across the talin molecule. These observations identify the first mechanically-gated anchoring protein for PKA, a new force-dependent binding partner for talin1, and thus a new mechanism for coupling cellular tension and signal transduction.

## Introduction

The cAMP-dependent protein kinase (Protein Kinase A; PKA) is the major receptor for the second messenger cAMP and is responsible for regulating myriad physiological and cellular processes. PKA is a heterotetrameric enzyme consisting of a dimer of regulatory (R) subunits that binds and sequesters two catalytic (C) subunits until cAMP binding to the R subunits causes C subunit release and activation (1, 2). PKA is ubiquitous and has hundreds of substrates associated with numerous distinct signaling pathways and cellular functions throughout the cell (3), leading to an abiding challenge and effort to understand how specificity in PKA signaling is achieved.

A significant literature establishes that, despite its ubiquity, PKA and its activity are highly localized within cells (4-6). While PKA C subunits were classically thought to freely diffuse from cAMP-bound R subunits (2), recent impactful work supports a more restrained radius of activity through either maintenance of an intact, catalytically-active tetrameric holoenzyme (7, 8) or restricted release and re-capture through membrane tethering of the C subunit (9, 10). Importantly, PKA activity is also localized through interaction (predominantly of type-II R subunits) with A-kinase anchoring proteins (AKAPs) that physically and functionally assign PKA to discrete subcellular niches (5, 11). AKAPs comprise a large, growing, and functionally diverse family of proteins with two common structural features: an amphipathic α-helix that mediates binding to R subunit dimers with nanomolar affinity and a complement of unique domains that specify distinct subcellular localization. In addition to anchoring PKA, AKAPs also often scaffold substrates and regulators of PKA as well as other signaling proteins (5, 11). Thus, AKAPs mediate the assembly and localization of discrete subcellular signaling nodes for PKA and allow this single kinase to participate in multiple, spatiotemporally distinct and dynamic cellular processes.

In this manner, AKAP-mediated localization of PKA activity has been shown to regulate various aspects of cell migration (12-14). For example, PKA activity is enriched in the leading edge in a manner that requires both anchoring and actomyosin contractility (15-17). PKA has also been shown to be both a complex regulator and effector of integrin-mediated adhesion to the extracellular matrix (ECM) (12, 14, 18-28), although the underlying molecular mechanisms are not fully understood. Recently (29), we reported the presence of PKA subunits, activity, and novel substrates within focal adhesions (FA), dynamic, multi-protein junctions formed between the cytoplasmic tails of ECM-bound integrins and the actin cytoskeleton (30, 31). Their position at this nexus means that FAs are subject to actomyosin-generated mechanical forces that alter the conformation of various FA components and thus regulate FA composition and dynamics (32-34). In addition to their structural role, FAs are solid-state signaling centers that scaffold kinases, phosphatases, and other enzymes to transmit and control the state of integrin-mediated cell adhesion and mechanosensing (30, 32-38). Central to this mechanosensitive machinery is the protein talin, which directly couples integrins to F-actin (39, 40). The C-terminus of talin is an extended rod comprised of 62 α-helices arranged in 13 modular bundles, R1-R13, that serve as force-dependent switch domains by opening and closing in response to small changes in contractility, thereby recruiting and displacing a myriad of interacting proteins enabling talin to serve as a mechanosensitive signaling hub (40). While many proteins have been identified that bind to the folded switch domains, only vinculin has been reported to bind to the unfolded domains, as its binding requires exposure of cryptic vinculin binding sites, comprising single amphipathic helices where the epitope is buried within the folded rod domain (41-44).

The functional connections between PKA and cell adhesion (12, 14), the presence of PKA subunits and activity within FAs (29), and the precept that AKAPs establish and co-localize with PKA activity microdomains (45) led us to search for potential AKAPs within FAs. In this study, we identify that PKA binds directly to an α-helix in talin that is cryptic and exposed only when the R9 domain unfolds, establishing talin as a mechanically-gated AKAP and PKA as the first mechanically-gated signaling partner for talin.

## Results

### Identification of PKA RIIα-binding proteins in FA

As a preliminary test for the presence of AKAPs in FA, whole-cell extracts (WCE) and isolated focal <u>a</u>dhesion <u>c</u>yto<u>s</u>keletal (FACS) fractions were analyzed using an RII overlay assay (46), a blotting method in which a purified recombinant V5 epitope-tagged docking/dimerization domain of PKA RIIα (RIIα-D/D) is used in place of a primary antibody, allowing detection of AKAP-RIIα interactions using an anti-V5 secondary antibody. Both whole cell extracts and isolated FACS fractions exhibited broad RIIα-binding activity over a wide range of molecular weights (*SI Appendix*, Fig.S1) that was significantly diminished by Ht31, an AKAP-derived peptide that binds to PKA R subunit D/D domains and thus competitively inhibits PKA-AKAP interactions (46), but not by the inactive control peptide Ht31p. These observations indicate the presence of AKAPs in FACS fractions.

To identify potential AKAPs within FA, we used proximity-dependent biotinylation catalyzed by a V5 epitope-tagged miniTurbo (mTb) biotin ligase (47) attached to the C-terminus of PKA RIIα. The mTb ligase biotinylates proteins in a ∼35 nm radius (48), so we hypothesized that RIIα-mTb would biotinylate AKAPs and proximal AKAP-associated proteins (Fig. 1*A*). Expression of V5 epitope-tagged RIIα-mTb resulted in robust protein biotinylation in a manner dependent on exogenous biotin (*SI Appendix*, Fig. S2*A*) and with a pattern very distinct from that catalyzed by ER-mTb, a well-characterized mTb targeted to the ER membrane (47). To increase experimental control of biotin labelling, we generated a stable line of U2OS cells in which the expression of V5-RIIα-mTb and consequent biotinylation are tightly controlled by a doxycycline-inducible promoter (Fig. 1*B, C*). Importantly, induced expression of V5-RIIα-mTb did not suppress bulk PKA activity (*SI Appendix*, Fig. S2*B*). Moreover, V5-RIIα-mTb, like endogenous RIIα subunits (29), was present in isolated FACS fractions (Fig. 1*D*) and catalyzed biotinylation of FACS proteins (Fig. 1*E*). To identify potential FA AKAPs, FACS fractions from RIIα-mTb-expressing cells (or uninduced cells, as a control) were collected and biotinylated proteins were isolated using streptavidin beads then characterized by LC-MS/MS-based proteomics (Fig. 1*F*). A total of 326 proteins were present either exclusively or at least five-fold higher in doxycycline-induced *vs* uninduced samples (Fig. 1*G*). This list was narrowed as shown in Fig. 1*G*. Briefly, we excluded proteins not present in the ‘meta-adhesome’ (a compilation of FA proteomic datasets (34, 49-52)), leaving 230 hits, and further refined this list by selecting proteins in the more exclusive ‘consensus core adhesome’ (34), comprising proteins common to all published FA protein lists. This list of 18 hits was then compared to a published list of *in silico* predicted AKAPs (53) generated by algorithmic scanning and scoring of protein sequences for homology to the degenerate PKA RIIα binding motif (see *SI Appendix*, Fig. S5). To increase stringency of this comparison, only proteins with a MAST (Motif Alignment and Search Tool, MEME Suite) score ≤ 50 (a cutoff that retains >95% of known AKAPs (53)) were considered. The only proteins surviving this vetting were talin1 and talin2, two isoforms of the key regulator of integrin-cytoskeletal coupling (40). Given that the number of talin1 peptides identified by LC-MS/MS greatly outnumbered that of talin2 (337 vs 2; *SI Appendix*, Dataset S1), we chose talin1 for initial investigation.

**Fig. 1.**
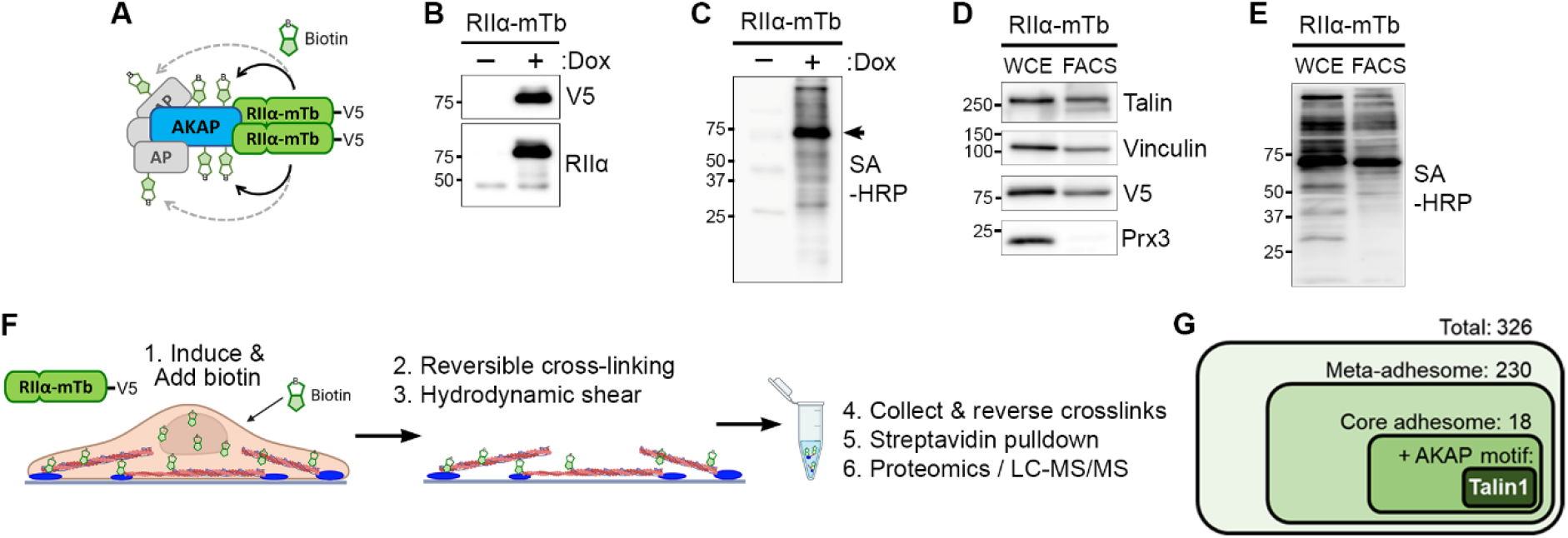
A proximity-dependent biotin labeling method reveals potential focal adhesion AKAPs. **(*A*)** Schematic of the theoretical approach, using the promiscuous biotin ligase miniTurbo (*mTb*) fused to PKA RIIα to biotinylate AKAPs and associated proteins (*AP*). **(*B, C*)** A stable line of U2OS cells expressing V5-tagged PKA RIIα-mTb under a doxycycline-inducible promoter were treated with doxycycline (*Dox*), labeled with biotin for 3 h, and lysates were analyzed by blotting with the indicated antibodies (*B*) or with streptavidin-conjugated HRP (*SA-HRP*; *C*). **(*D, E*)** Whole cell extracts (*WCE*) or focal adhesion/cytoskeleton fractions (*FACS*) were made from Dox-induced cells and analyzed by immunoblotting with the indicated antibodies or with SA-HRP. **(*F*)** Schematic of the FA AKAP screening protocol. Biotinylation of RIIα binding proteins after induction of PKA RIIα-mTb and addition of biotin (1) is followed by isolation of FA proteins *via* reversible cross-linking (2) and hydrodynamic unroofing of cells (3). The remaining FACS fractions are collected and cross-linking is reversed (4), then biotinylated proteins are captured on immobilized avidin (5) then trypsinized and analyzed by LC-MS/MS-based proteomics (6). **(*G*)** Bioinformatic pipeline for vetting candidate FA AKAPs. Hits from proteomic analysis were initially screened for inclusion in a combined ‘meta-adhesome’ then for inclusion in the consensus core adhesome, and for the presence of a high-scoring, *in silico*-predicted potential AKAP motif. The only hit to survive this vetting was talin1.

### Talin1 is a novel AKAP

Talin1 is a crucial FA component, binding to and activating integrins and coupling them directly to cytoskeletal actin (39, 40). Talin1 also contributes to integrin-mediated adhesion through interaction with myriad FA and signaling proteins that instigate FA assembly and regulate adhesive signaling and mechanosensing (39, 40). To determine whether talin1 is indeed an AKAP, we directly assessed talin-PKA interaction. Talin immunoprecipitated from biotin-labeled, V5-RIIα-mTb-expressing cells is detectable by streptavidin-HRP (Fig. 2*A*), confirming that talin is directly biotinylated by RIIα-mTb and not present in the dataset due to indirect interaction with a biotinylated intermediary protein. Expression of an RIIα-mTb with double point-mutations (I3S, I5S) known to disrupt RII-AKAP interactions (54, 55) significantly reduced the amount of both AKAP79 (a well-characterized AKAP (56) used as a control) and talin in streptavidin pull-downs compared to WT RIIα-mTb levels (Fig. 2*B*). Furthermore, both AKAP79 and talin co-immunoprecipitated with PKA RIIα-mTb under cross-linking conditions (Fig. 2*C*). Taken together, these results support a close molecular interaction between talin and PKA RIIα but do not directly demonstrate that the interaction is direct – a *sine qua non* requirement for classification as an AKAP.

**Fig. 2.**
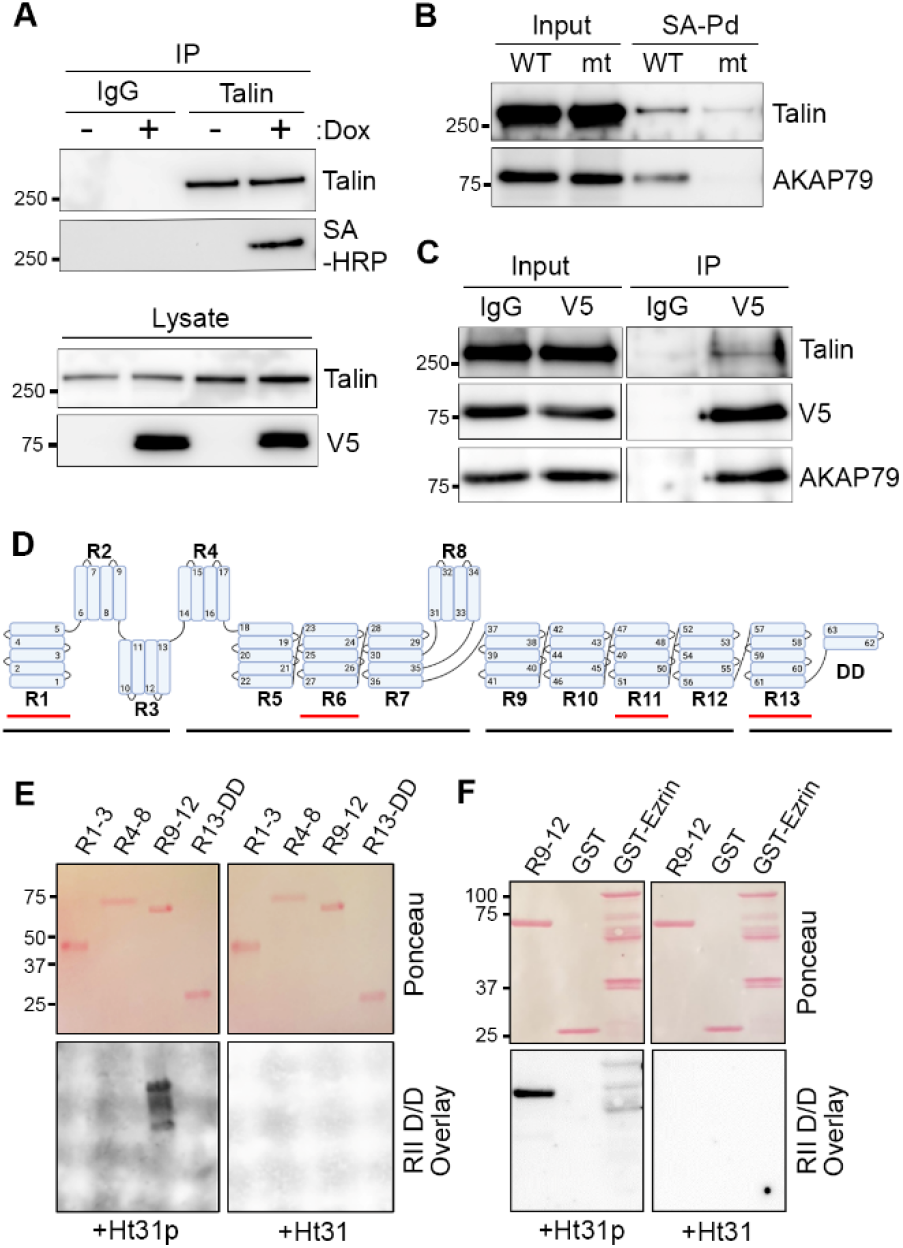
Talin is an AKAP. **(*A*)** Cells were treated, where indicated, with Dox to induce V5-RIIα-mTb expression and labelled with biotin for 3 h. Lysates were directly blotted for talin or RIIα-mTb, or were immunoprecipitated with control IgG or anti-talin antibody then blotted with anti-talin antibody or streptavidin-HRP. **(*B*)** Cells induced to express wild-type (*WT*) RIIα-mTb or a point mutant deficient for AKAP binding (*mt*) were labelled with biotin. Lysates (*Input*) or streptavidin pull-downs (*SA-Pd*) were analyzed by immunoblotting for talin or AKAP79. **(*C*)** Lysates and control (*IgG*) or anti-V5 immunoprecipitates of DTBP-crosslinked, V5-RIIα-mTb-expressing cells were analyzed by immunoblotting with the indicated antibodies. **(*D*)** Schematic of the talin1 rod domain, indicating the 13 rod (R) domains, R1-13 and the four fragments, R1-3, R4-8, R9-12 and R13-DD (*black underline*) used below. Each fragment contains an *in-silico*-predicted RII-binding domain (*red underline*). **(*E*)** The fragments indicated in (*D*) were expressed recombinantly, separated by SDS-PAGE and transferred to membranes which were analyzed for equal loading (*Ponceau*). After destaining, membranes were analyzed for direct PKA interaction by overlay with the V5-tagged RIIα-dimerization/docking (*RII D/D*) domain in the presence of either a competitive peptide that blocks AKAP binding (*Ht31*) or a non-blocking negative control peptide (*Ht31p*). **(*F*)** Talin1 R9-12 fragment, GST alone and GST-tagged ezrin were analyzed by RII D/D overlay as described in (*E*).

Talin1 is a large (∼270 kDa) protein comprising an N-terminal globular head domain connected to an extensible rod domain comprising 62 helices arranged in 13 helical bundle domains (R1–R13) with a helical dimerization domain (DD) at the C-terminus (39, 57). *In silico* AKAP prediction software identified four of these 62 amphipathic helices (Fig 2*D*) as putative PKA RIIα binding sites (53). To test whether talin interacts directly with PKA RIIα, constructs that divide the rod into 4 fragments – R1-3, R4-8, R9-12 and R13-DD (Fig 2*D*) – for recombinant protein expression were used (58). Fortuitously, each of these fragments contains one of the four *in silico*-predicted RIIα binding sites (Fig. 2*D*), so the purified fragments were analyzed by PKA RIIα overlay assay. Importantly, the denaturation and *in situ* refolding steps in this assay promotes refolding of individual α-helices but not of higher-order tertiary structures such as helical bundles (59-61). Strikingly, only the fragment comprising talin1 rod domains R9-12 bound directly to both the PKA RIIα D/D domain (Fig. 2*E*) and to purified, full-length PKA RIIα (*SI Appendix*, Fig. S3). This interaction could be specifically inhibited by the Ht31 inhibitor peptide indicating a canonical AKAP interaction. Furthermore, the interaction of talin1 R9-12 with PKA RIIα was significantly stronger than that of the known AKAP, ezrin (Fig. 2F), a membrane-associated cytoskeletal linker (62, 63). Taken together, these data indicate that talin1 directly binds PKA RIIα and establishes talin1 as a new AKAP.

### The R9 and R10 domains of talin1 are required, but neither is sufficient, for PKA RIIα Binding

The *in silico*-predicted PKA RIIα binding motif within the R9-R12 fragment is helix 50 (h50) in the 5-helix R11 domain (*SI Appendix*, Fig. S4), which shows significant homology to the canonical amphipathic helical motif for AKAPs (*SI Appendix*, Fig. S5). To determine whether talin h50 mediates binding to PKA, we mutated a key residue on the hydrophobic face of the helix, V2087, which would be predicted to abolish interaction with PKA RIIα (46). Specifically, we generated three mutant talin1 R9-12 fragments targeting V2087 (V2087P, V2087S, and V2087A). V2087P introduces a helix-breaking proline (a mutation analogous to the one used in the inactive Ht31p control peptide (46)), while V2087S interrupts the hydrophobic surface of h50 and the conservative V2087A replaces valine with the smallest hydrophobic residue alanine. Surprisingly, all mutant fragments, including V2087P, bound PKA RIIα D/D in overlay assays (*SI Appendix*, Fig. S4), strongly suggesting that h50, despite predictions and homology, is not the PKA RII binding site in talin.

We next reverted to an empirical approach to map the RII binding site on talin1, performing overlay assays with serial truncations of the RIIα-binding R9-12 fragment (Fig. 3*A*). Removal of R12 and R11 did not disrupt talin-RIIα interaction, confirming that h50 is not the PKA binding site. Furthermore, while R9-10 still bound RIIα D/D, an isolated R9 domain did not (Fig. 3*A*), suggesting that R10 is essential for PKA RIIα binding. To confirm this, we performed additional overlay assays with R9-12 (ΔR10) and R10 alone (Fig. 3*B*). As expected, R9-12 (ΔR10) did not bind PKA RIIα D/D, reinforcing the hypothesized requirement for R10. Surprisingly, however, the R10 domain showed only very weak RIIα binding on its own (Fig. 3*B*). Together, these data indicate that the R9-10 domains are required for efficient binding to PKA and that neither R9 nor R10 alone is sufficient for binding.

**Fig. 3.**
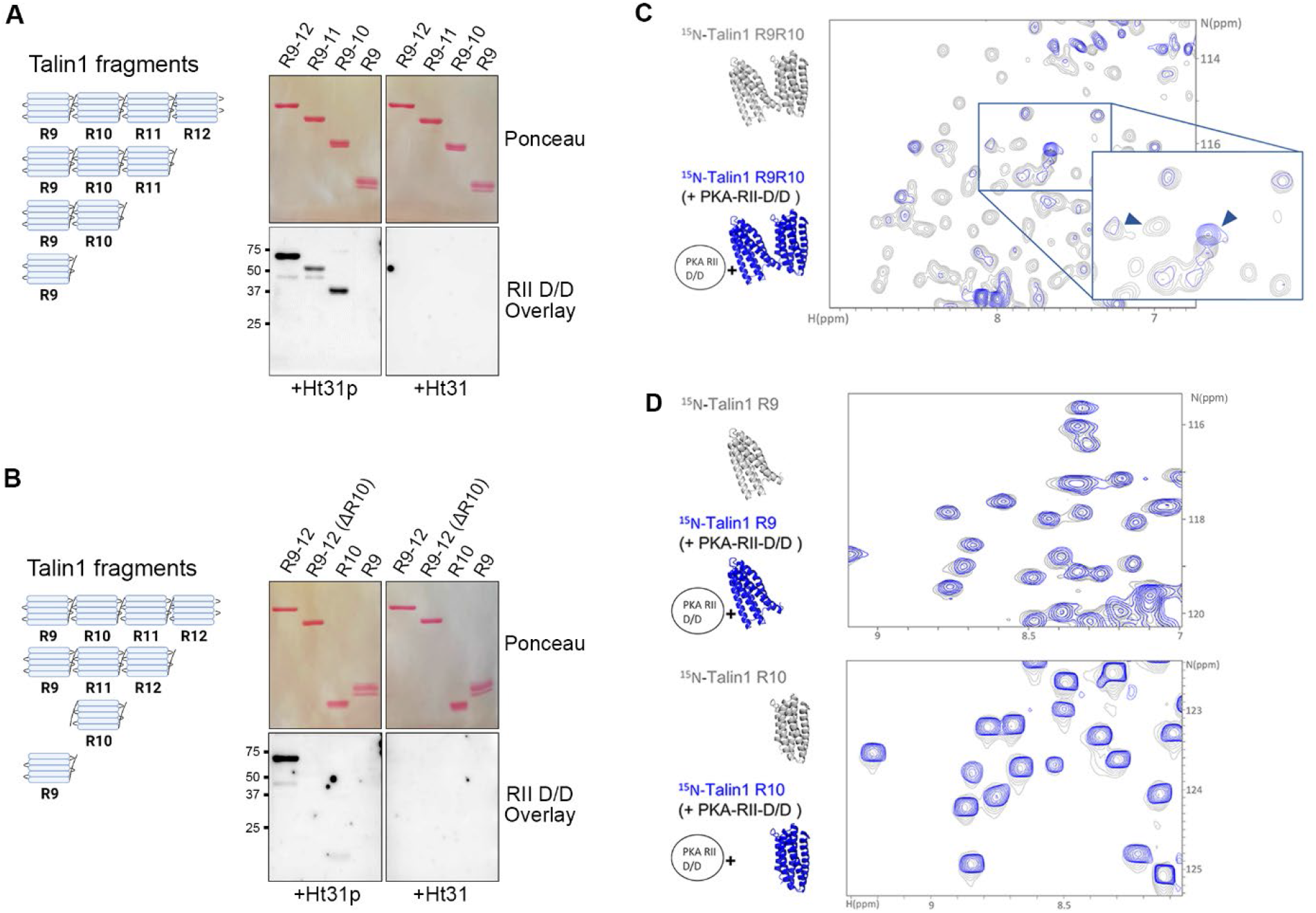
The R9 and R10 domains of talin1 are required, but neither is sufficient, for PKA RII D/D binding. **(*A, B*)** Recombinant talin R9-12 and various R-subdomain truncation, deletion, or isolation fragments (*left*) were analyzed by RII D/D overlay (*right*) in the presence of either control Ht31p or AKAP-blocking Ht31 inhibitor peptides. **(*C*)** ^1^H,^15^N HSQC spectra of ^15^N-labeled talin R9-10 domains in the absence (*grey*) or presence (*blue*) of PKA RII D/D domain at a ratio of 1:5. Arrows indicate spectral peaks that disappear or shift in the presence of RII D/D. **(*D*)** ^1^H,^15^N HSQC spectra of individual ^15^N-labeled talin R9 (*top*) or R10 (*bottom*) domains in the absence (*grey*) or presence (*blue*) of PKA RII D/D at a ratio of 1:3.

### NMR spectroscopy reveals that talin R9 contains the RIIα binding site

The challenge of interpreting these RII overlay results prompted us to adopt a different methodology to assess the structural basis of the talin-PKA interaction. Thus, we used protein NMR spectroscopy – specifically, ^1^H,^15^N TROSY-HSQC experiments. HSQC spectra provide a powerful way to evaluate the structure of a protein, as each backbone amide gives rise to a discrete peak. When an unlabeled binding partner is added, changes to the spectral peaks can be detected. As reported previously, the NMR spectrum of talin1 R9-R10 shows good peak dispersion (57), and the spectra of the individual R9 and R10 domains overlay well indicating only limited interaction between the two domains (Fig. 3*C* and (57)). Addition of unlabeled PKA RIIα D/D to ^15^N-labeled R9-10 talin fragment confirmed a direct interaction of PKA to talin1 R9-10, as evidenced by significant changes in the talin spectra (Fig. 3*C*). A similar experiment with talin2 R9-R10 demonstrated that both talins can bind PKA (*SI Appendix*, Fig. S6). The nature of the spectral changes upon talin-PKA interaction was unexpected, however, as it showed a dramatic loss of a subset of the signals, which were predominantly from the talin R9 domain, with the signals from the R10 domain largely unaffected (Fig. 3*C* and *SI Appendix*, Fig. S7). Specifically, addition of PKA RIIα D/D to talin R9-10 caused the signals from R9 to broaden and decrease in intensity (Fig. 3*C* and *SI Appendix*, Fig. S7*A*). This suggests that RIIα binding to R9-10 causes R9 to unfold and adopt a molten globule-like state, with broadened peaks indicative of an unfolded bundle but lacking the strong signal intensity of a fully-extended polypeptide. In contrast, PKA RIIα D/D had minimal effect on the NMR spectra of individual R9 or R10 fragments, with only very small chemical shift changes seen in R9 indicative of a very weak interaction but no alteration in structure (Fig. 3*D*), consistent with the inability of individual R9 or R10 to bind PKA in overlay assays (Fig. 3*B*). Together this NMR analysis indicates that the PKA RIIα D/D binds to R9-R10 primarily on R9, and is able to unfold the R9 domain but, intriguingly, only when its attached to R10.

### Helix41 in talin R9 domain is the PKA RIIα binding site, but it requires the R9-R10 linker and part of Helix42 from R10

AKAPs canonically use the hydrophobic face of an amphipathic helix to bind the RII dimer, and talin helices are arranged in bundles with the hydrophobic surfaces towards the center. Thus, we hypothesized that the observed R9 unfolding is due to the RII-binding motif being on residues of a helix that are buried within the folded R9 5-helix bundle and thus high-affinity PKA binding requires unfolding of the talin1 R9 domain to expose this motif. The paradigm of talin helix bundles unfolding in order to bind ligands is established *via* the well-studied interaction between vinculin and the 11 talin helices that are vinculin-binding sites (VBS) (39, 40). However, to date, vinculin is the only protein identified that binds to open talin bundles. Furthermore, vinculin binding does not require an adjacent domain being present to bind, unlike the currently observed requirements for PKA binding. Therefore, we next set out to identify how R10 was contributing to R9 unfolding and binding to PKA RII D/D. When we first resolved the domain boundaries of R10, we generated a series of constructs of R10 (64), including one 6-helix fragment – helix41-R10 – comprising the last helix of R9 and the five helices of R10 (helices 41-46), so we thought to test whether this fragment was sufficient to bind PKA RIIα subunits. In the spectra of h41-R10, the signals corresponding to h41 are readily identifiable by overlaying the spectra of R10 alone (*SI Appendix*, Fig. S8 and (64)). As the isolated h41 is unbundled in this construct, it is unstructured, has only helical propensity and, as a result, its NMR peaks form a cluster of sharp signals with poor dispersion (*SI Appendix*, Fig. S8 and (64)), making it particularly amenable for visualizing shifts upon protein-protein interactions. Thus, ^15^N-labeled talin1 h41-R10 fragment was analyzed by NMR in the absence and presence of PKA RIIα D/D (Fig. 4*A*). Upon addition of PKA RIIα D/D, chemical shift changes were visible, indicating binding of PKA to h41-R10. Moreover, the shifts were predominantly in the intense peaks that correspond to unbundled h41, and the signals both shifted and became less sharp as the helical conformation is stabilized upon binding (Fig. 4*A*).

**Fig. 4.**
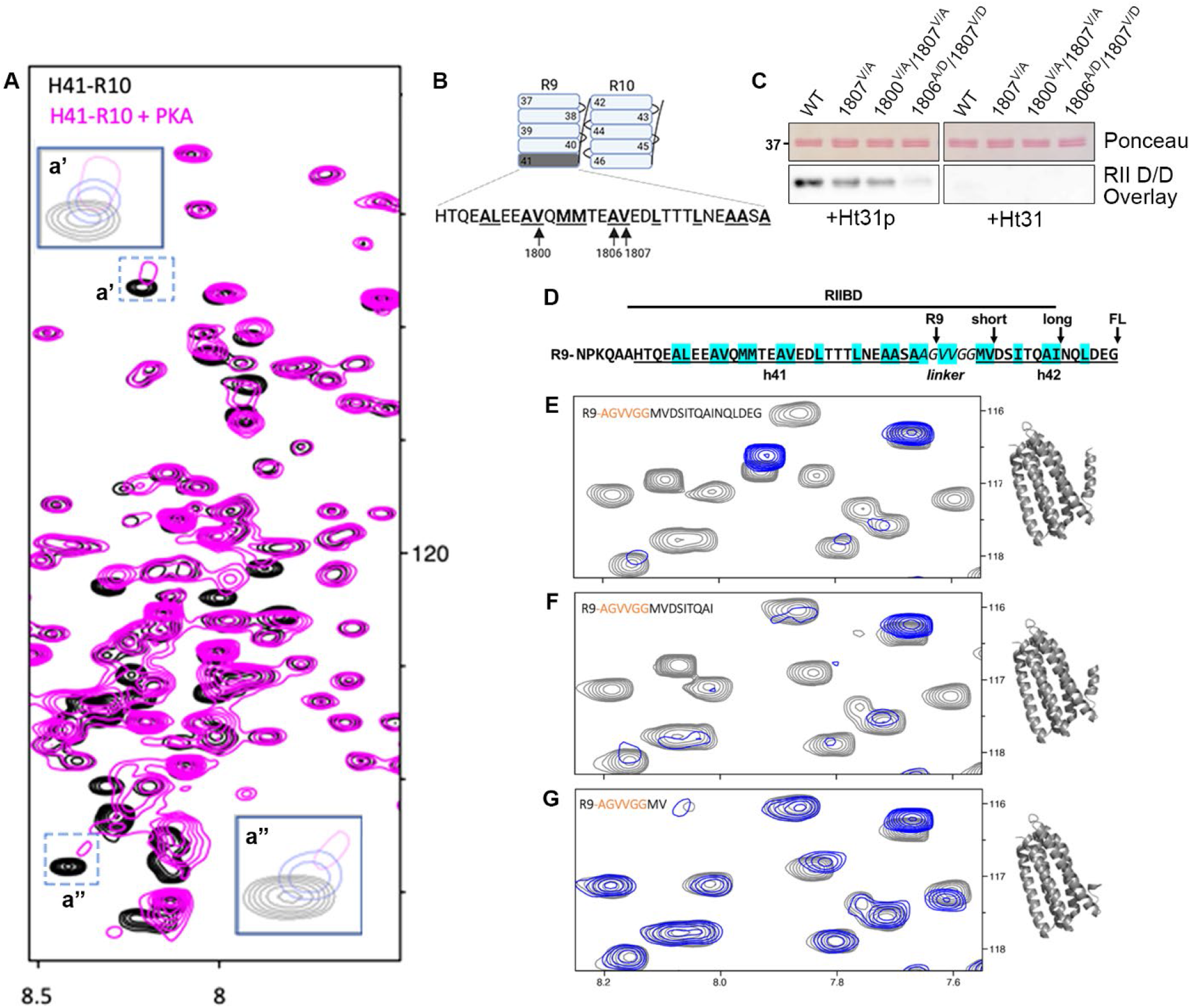
Helix41, the R9-R10 linker, and part of helix42 together are required for PKA RII binding. **(*A*)** ^1^H,^15^N-HSQC spectra of ^15^N-labeled talin1 helix41-R10 (h41-R10; residues 1785-1973) in the absence (*black*) or presence (*magenta*) of PKA RII D/D domain at a ratio of 1:5. The intense peaks from h41 are the ones that shift. Two insets (*a’, a’’*) show magnification of regions (indicated by dotted outlines) with peaks that shift on addition of increasing amounts of PKA RII D/D (1:0, 1:3. 1:5 in black, blue, and magenta, respectively). **(*B, C*)** Three single or double point mutants, targeting selected hydrophobic residues within helix41 (*B*), were generated and analyzed by PKA RII D/D domain overlay (*C*). **(*D*)** Amino acid sequence of h41-linker-h42. Underlining indicates amino acids in helices, italics indicate the linker, and highlighting indicates hydrophobic residues. The terminus of R9-h42 (*FL*), long and short h42 truncations, and the R9 domain-only construct are indicated with arrows (*RIIBD* = RII binding domain). **(*E-G*)** ^1^H,^15^N-HSQC spectra of ^15^N-labeled talin R9-h42 and truncated (*long, short*) fragments indicated in the absence (*grey*) or presence (*blue*) of PKA RII D/D domain fragments at a ratio of 1:3.

Alignment of the sequence of h41 with the PKA RII-binding motifs of known AKAPs shows significant similarities (*SI Appendix*, Fig. S5), most notably the semi-regular spacing of aliphatic residues that form the requisite hydrophobic face on the helix. This alignment also reveals some significant deviations, most notably a pair of methionine residues in a position typically occupied by smaller aliphatic amino acids (Fig 4*B* and *SI Appendix*, Fig. S5 (5, 46, 54)) – a deviation often associated with decreased affinity for RII subunits (46, 65) and likely responsible for the absence of h41 from the *in silico*-predicted talin1 AKAP motifs (53). Given these considerations, along with the unique requirement of unfolding of R9 to enable h41 to bind RIIα subunits, we endeavored to confirm the principal importance of h41 as the primary RIIα-binding interface using targeted point mutations (Fig. 4*B & C*). Overlay assays revealed that two mutants with conservative substitutions (V1807A and V1800A/V1807A) still bound RIIα D/D, albeit at a significantly reduced level, while a double mutant with more radical substitutions that disrupts the hydrophobic face of h41 (A1806D/V1807D) exhibited markedly reduced binding (Fig 4*C*), confirming the importance of h41 for PKA RII binding. These results unequivocally establish h41 as the PKA RIIα-binding site in talin1.

Together, the preceding observations demonstrate that h41 is the RIIα binding site and is available for PKA binding when it is ‘free’ or unbundled, as in h41-R10 (Fig. 4*A*). However, in the context of the complete R9 domain bundle, the site is ‘cryptic’ and requires unfolding of R9 and exposure of the hydrophobic face of h41 for RIIα access (Fig. 3*B-D*). In these experiments, this unfolding requires R10. Therefore, we next wanted to determine how much of the R10 domain is required to unfold R9 and allow PKA to bind. To do this we analyzed an ^15^N-labeled talin1 fragment comprising R9-h42 (R9 through to the first full helix of R10; Fig. 4*D*) and found that addition of PKA RIIα D/D to R9-h42 elicited clear, striking changes in R9 spectral peaks, indicating that the addition of h42 alone was sufficient to allow PKA to unfold and bind to R9 (Fig. 4*E*). Thus, both h41-R10 and R9-h42 are able to bind PKA – one because the PKA binding site is constitutively exposed (h41-R10) and the other because PKA is able to unfold R9 to expose it (R9-h42).

To further define how much of the sequence beyond R9/h41 is required to allow PKA to unfold R9 and bind, we analyzed a series of fragments of R9-h42 (Fig. 4A & 4*E*), with decreasing lengths of the R9-R10 linker and h42 (Fig. 4*D*), approaching the R9 domain alone which is not unfolded by PKA (Fig. 3*D*). Thus, R9 with 13 additional residues (spanning the entire R9-R10 linker and first ten of the 16 residues of h42; *R9-h42-long*) unfolded and bound to PKA RIIα D/D as efficiently as the full-length R9-h42 (Fig. 4*F*). However, R9 with only 6 additional residues (spanning the entire R9-R10 linker and first two residues of h42; *R9-h42-short*) showed only weak binding to PKA RIIα D/D and no unfolding of R9 (Fig. 4*G*). Recall that removal of these six residues generates R9 alone which, as shown above, does not bind (Fig. 3*D*). These data indicate that the interaction of PKA RIIα to talin1 is mediated by h41 in R9, but in a manner that requires the R9-R10 linker and a small portion of h42 in R10 for unfolding, exposure, and strong binding. Based on these cumulative data, we assign the region of amino acids 1785-1835 (h41-linker-h42-long) as the talin1 PKA RII binding domain (RIIBD; Fig. 4*D* and *SI Appendix*, Table S1). Interestingly, the first six residues of the h42 sequence in R10, that are required for PKA binding to R9, are not a well-defined part of the helix in the structures containing R10 (64) which likely explains why PKA binding does not unfold, nor require unfolding of, R10.

### Mechanical force across talin regulates PKA RIIα Binding

At the nexus of ECM-bound integrins and the actomyosin cytoskeleton, talin1 functions not only as a linker but also as a mechanosensitive scaffold, with force-dependent unfolding of its rod domains controlling interactions with various binding partners (39, 40, 43). The most well-studied example of this mechanosensitivity is the binding of talin to vinculin, with elegant structural and biophysical studies establishing that this interaction requires unfolding of talin rod bundles containing cryptic VBS and demonstrating that this conformation could be attained *in vitro* by application of mechanical force across relevant talin rod domain fragments (41, 42, 66, 67). Given that PKA binding to talin1 requires the talin R9 domain to unfold, as described above, it has the potential to be similarly regulated by mechanical force. This would be consistent with the current observation that, while talin1 is one of the most abundant proteins biotinylated by RIIα-mTb, co-immunoprecipitation of talin1 with RIIα-mTb occurred only under crosslinking conditions (Fig. 2*C*).

To directly test this hypothesis, we used magnetic tweezers to immobilize and mechanically stretch the talin1 R9-12 fragment, as described previously (68), to generate force-extension curves before and after addition of PKA RIIα D/D. For each force cycle, a linearly increasing force from 1.5 to ∼30 pN is applied to the protein tether at a constant loading rate to unfold the domains, then the applied force is reduced to ∼1.5 pN to allow refolding. During force loading, the height of the end-attached superparamagnetic microbead was recorded at a nanometer resolution in real time (69, 70). In the absence of PKA RIIα D/D, R9–12 shows four distinct unfolding steps in the force range of ∼10–25 pN (Fig. 5*A*), with each extension of ∼35–50 nm corresponding to the unfolding of a rod domain bundle as seen previously (68). These results confirm that, in the absence of other proteins, all the four domains unfold rapidly once the applied force exceeds the mechanical stability of that domain, and that they each faithfully refold at low forces.

**Fig. 5.**
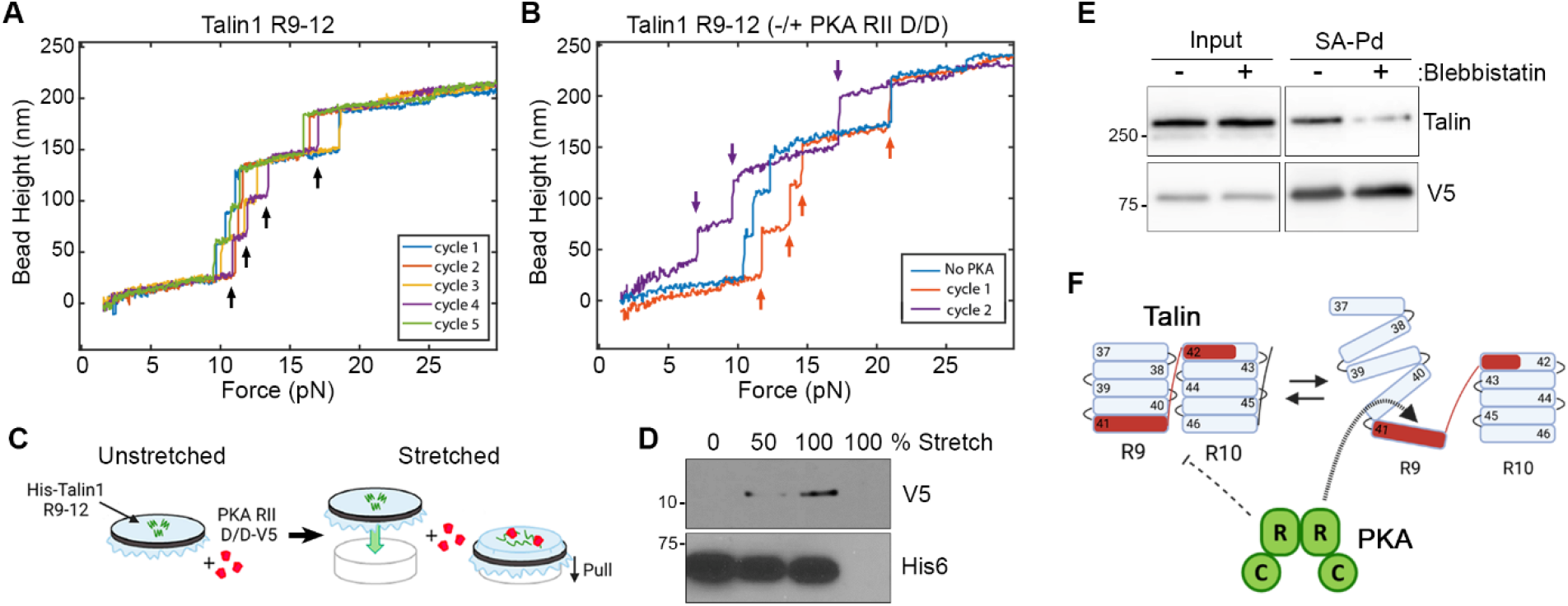
Mechanical gating of talin-PKA interaction. **(*A, B*)** Talin1 R9-12 was tethered between a coverslip and a paramagnetic bead and magnetic tweezers were used to generate unfolding force *vs* bead height (*i*.*e*., extension) curves. (*A*) Multiple extension cycles of a single talin R9-12 tether alone, demonstrating four characteristic extensions corresponding to the unfolding of each of the four R domains (*black arrows*). (*B*) Iterative cycles of a single talin R9-12 tether before (*blue*) and after (*orange, purple*) addition of 100 nM PKA RIIα D/D domain. The characteristic four extensions persist for one extension cycle after addition of PKA D/D (*orange arrows*) but the number of unflolding steps is reduced to three (*purple arrows*) in the subsequent cycle, indicating unfolding-dependent binding of PKA RIIα D/D which results in inhibition of the refolding of one domain. **(*C*)** Schematic of the immobilized protein extension (IPE) assay. A bait protein is adsorbed onto a taught flexible silicon sheet which is left taught or is stretched to varying extents before incubating with a prey protein. Bound proteins are collected and analyzed by SDS-PAGE and immunoblotting. **(*D*)** IPE assays of uncoated or His_6_(R9-12)-coated sheets that were unstretched (*0%*) or stretched by the indicated amounts (as % increase in surface area) before incubation with V5-tagged PKA RIIα D/D domain. Bound proteins were collected and immunoblotted with the indicated anti-tag antibodies. **(*E*)** V5-RIIα-mTb expressing cells were labeled with biotin in the absence or presence of 25 μM of blebbistatin. Lysates or biotinylated proteins isolated using streptavidin bead pull-down (*SA-Pd*) were analyzed by immunoblotting with the indicated antibodies. **(*F*)** A hypothetical model of mechanically-gated talin-PKA RII interaction where unfolding of the R9 domain leads to exposure of the talin AKAP and binding of PKA.

Importantly, this characteristic force-bead height profile is significantly altered after addition of PKA RIIα D/D (Fig. 5*B* and *SI Appendix*, Fig. S9). Specifically, in the first cycle in the presence of 100 nM PKA RIIα D/D, the characteristic four unfolding steps of R9-12 are still present, indicating that PKA addition alone was not sufficient to unfold the domains. However, within 1-2 force loading cycles, the number of unfolding steps decreased from four to three, and correspondingly the bead height increased at forces below 10 pN. Moreover, the three unfolding steps persisted in subsequent force-loading cycles (Fig. 5*B* and *SI Appendix*, Fig. S9), indicating that one domain was unable to refold. Together, these data indicate an unfolding-dependent binding of PKA RIIα D/D to talin R9-12 and subsequent inhibition of the refolding of one domain.

We noticed with some tethers that increased numbers of force loading cycles (typically 3-4 cycles) in the presence of RIIα D/D led to a reduction in the maximum extension length of R9-12 at the tested maximal force (*SI Appendix*, Fig. S10), suggesting the possibility of an additional interaction between PKA RIIα D/D and unfolded α-helices in R9-R12. However, such an additional interaction was not evident in our biochemical assays. We note that maintaining R9 in an unfolded state exposes two buried cysteines (C1661 and C1671), and even though the experiments were done in the presence of reducing agent (2 mM TCEP), we cannot exclude that this is not a result of looping via a disulfide bond that is only able to form because the R9 is prevented from refolding. Nonetheless, the reduced number of unfolding steps persisted through multiple force-loading cycles even as the tether length was reduced (Fig. 5*B* and *SI Appendix*, Figs. S9 and S10). Taken together, these observations unequivocally demonstrate the regulation of talin-PKA interaction by force-dependent changes in talin conformation, and show that PKA RIIα D/D stabilizes the unfolded conformation of one of the domains through multiple cycles of force-loading.

We next corroborated the observations from these single-molecule biophysical experiments using a previously described immobilized protein extension (IPE) assay (71) in which the talin1 R9-12 fragment was adsorbed onto a silicon membrane before being incubated with RIIα-D/D under unstretched or stretched conditions (Fig. 5*B*). While only qualitative, this assay clearly showed that RIIα-D/D bound to immobilized talin1 R9-12 in proportion to the level of membrane stretch with minimal binding in the absence of stretch (Fig. 5*C*), further demonstrating direct mechanical regulation of the talin1-PKA interaction. Finally, we sought to further support these observations in cells by altering actomyosin contractility. We inhibited actomyosin contractility, using blebbistatin, in cells expressing the RIIα-mTb biotin ligase then assessed the amount of talin recovered in streptavidin pull-downs. While the level of auto-biotinylated RIIα-mTb remained unchanged, indicating no contractility-dependent changes in ligase activity, the level of talin in the blebbistatin-treated sample was significantly reduced (Fig. 5*D*). These data are consistent with a blebbistatin-induced reduction of contractility leading to decreased mechanical unfolding of talin1 (72) and reduction in talin1 biotinylation through decreased tension-dependent binding of PKA RIIα-mTb. Collectively, the observations in this report establish that talin is an anchoring protein for PKA and that the talin-PKA interaction is regulated by conformational changes in talin that are controlled by mechanical force.

## Discussion

It is increasingly well established that cellular signal transduction events are most often conducted through multi-protein scaffolds that consolidate, localize, and specify signaling inputs and outputs (5, 73). There is increasing recognition that proteins involved in cell-ECM adhesion are ideally positioned to convert mechanical force into altered biochemistry (74, 75). This report bridges these two important fields by identifying talin, the archetypal FA protein, as a new anchoring protein for PKA and the first mechanically-gated AKAP to be described. In so doing, it also establishes PKA, a pleiotropic kinase with myriad cellular targets, as a new force-dependent binding partner and signal transducer for talin1. Taken together, these observations form the foundation for a novel mechanotransduction pathway that utilizes force-dependent changes in protein conformation to establish a new, solid-state signaling complex that couples cellular tension to cellular communication.

Given the promiscuity of PKA enzymatic activity and the number and diversity of substrates, mechanisms have evolved to focus or target PKA function to enhance signaling fidelity and specificity. The best characterized mechanism for this is through AKAP-mediated changes in PKA localization, thereby promoting or sequestering PKA activity to specific subcellular niches, substrates, and events (5, 11, 14). The diversity of mammalian AKAP complexes, which differ not only in their localization but also in their composition and dynamics, produces a wide range of distinct signalosomes, each producing highly regulated and specific PKA outputs. Thus, identifying new AKAPs increases our understanding not only of how PKA contributes to specific cellular events but also of how signaling is controlled at the subcellular scale. In this way, the current demonstration of talin as an AKAP provides an important new niche for PKA signaling, while the demonstration that the PKA-talin interaction is regulated by mechanical force across talin establishes an important new type of mechano-regulated PKA anchoring.

The observations presented here are the first to show direct, mechanical gating of PKA anchoring. Indeed, the requirement for conformational rearrangement for AKAP-PKA interaction is not common – nearly all knowns AKAPs are reported to interact with PKA R subunits constitutively (46, 76, 77). To our knowledge, the only other exception to this is ezrin, a membrane-cytoskeleton adapter protein whose phosphorylation-dependent switch between open and closed conformations controls its binding to PKA type-I, but not RIIα, subunits (62, 78). The PKA binding site in talin identified here is similarly cryptic, being occluded when R9 is in its closed conformation and available for binding only when the domain is opened – by denaturation and partial refolding (as in the overlay assays), high stoichiometric excess of PKA D/D domain (as in the NMR assays), or mechanical extension through applied force. This requirement for conformation and/or mechanical gating, combined with the fact that the PKA binding domain in h41 of talin has significant divergence from canonical RIIα binding motifs (Fig. 4*D*; *SI Appendix*, Fig S5), suggests that the ‘fit’ between the RIIα dimer and a given AKAP can be modified and enhanced by active conformational changes within the AKAP itself. This, in turn, underscores the novelty of the PKA-talin complex and importance of determining its structural detail at higher resolution – a pursuit currently underway. It further suggests the possibility of additional, ‘semi-canonical’ AKAPs with divergent binding motifs that would not be readily predicted by *in silico* screening or algorithms. From these considerations, it is intriguing to posit the possibility of similar mechanically-regulated PKA anchors in other dynamic, tension-bearing cellular structures such as cell-cell junctions and kinetochores.

Conversely, the mechanically-regulated interaction between PKA and talin has important consequences for our understanding of talin biology as well. Talin is known to bind other enzymes (*e*.*g*. focal adhesion kinase, phosphatidylinositol 4-phosphate 5-kinase type I-γ, cyclin-dependent kinase-1 and alpha tubulin acetyltransferase 1 (39, 58, 79)) but the talin-PKA is noteworthy because it represents the first report of an enzyme that binds to a mechanically-opened rod domain. Prior to this discovery, the only other protein known to bind to the force-dependent open conformation of talin rod domains was vinculin, another core actin-binding FA protein and mechanotransducer (39, 41, 80, 81). Indeed, there are important parallels between the well-known, mechanically-regulated talin-vinculin interaction and the newly described talin-PKA interaction. Each of the eleven vinculin-binding sites (VBS) in talin consists of a string of hydrophobic residues along the face of an individual helix, which are buried within the folded rod domains. In the case of the talin-vinculin interaction, force across the talin rod exposes one or more of these sites, and the subsequent force-dependent, talin-mediated recruitment of vinculin to FAs is crucial for strengthening the link between integrins and F-actin (39, 80-82). As only 11 of the 61 helices that form the 13 talin rod domains have been reported to bind vinculin once exposed, the other 50 helices might also be expected to bind other ligands, yet evidence for this has been lacking until now. Therefore, the PKA-talin interaction confirms a new paradigm for talin signaling whereby signaling molecules and enzymes can be directly recruited to the exposed residues of open talin switch domains.

While not likely to contribute to architectural reinforcement directly, it is quite likely that recruitment of PKA to talin may, at the most immediate or localized level, promote the phosphorylation of talin itself and/or some of the myriad talin-associated proteins (39, 40, 83). Evidence for the former comes from our own preliminary experiments showing direct phosphorylation of purified talin1 R4-8 and R9-12 fragments and increased talin1 phosphorylation, recognized by an anti-phospho-PKA substrate antibody, in cells treated with treated with forskolin and IBMX which increase cAMP and activate PKA (*SI Appendix*, Fig. S11). More circumstantial, but nonetheless supportive, evidence comes from the presence of predicted PKA phosphorylation sites in the sequence of several talin-associated proteins (using PhosphoSitePlus, Phosida, and dbPAF; also (84)) and the presence of talin and several talin-associated proteins in unbiased screens for PKA substrates (85-91).

An important direction for future efforts involves delineating the cellular regulation and function of the talin-PKA complex. Talin is autoinhibited by an interaction between the folded R9 domain and the F3 domain in the talin head; as F3 contains the principal integrin binding site of talin, this interaction must be released to allow binding to actin and integrin and recruitment to FAs (39, 40, 83). We have shown that the major constituent of the PKA binding site in talin is h41 of R9, which therefore suggests that the talin-PKA interaction is most likely to occur at sites of talin activation and where R9 is unfolded. Similarly, PKA binding to talin might limit R9 refolding maintaining talin in an active conformation, and so activate and sustain adhesion assembly. Direct analysis of the force borne by talin in adhesive complexes in live cells, using a mechanically-responsive talin biosensor, established that talin is under higher tension in peripheral FAs compared to more central FAs and/or fibrillar adhesions, and even showed heterogenous force-loading within individual FAs (72, 92). In recent work, we have reported that PKA is active and dynamic within sub-regions of FA (29). Furthermore, as the opening of talin rod domains introduces 40-50 nm extension in the length of the talin molecule (93), the anchoring of PKA could be dynamically moved within each adhesion as other talin rod domains open and close. It will be an important and informative challenge to ascertain the extent to which talin-mediated anchoring is required for localized PKA activity and to further determine the precise subset of adhesion complexes (or sub-regions within individual adhesion complexes) in which the talin-PKA interaction occurs.

Finally, it will also be important to identify the physiological function of (*i*.*e*., the consequences of specific loss of) talin-mediated PKA anchoring for cell adhesion and motility. This effort will require the generation of discrete talin point mutants that prevent RIIα binding without altering any other talin function (*e*.*g*., talin autoinhibition, folding/refolding; other protein-protein interactions). Although we show here it is possible to disrupt the talin-PKA interaction using a canonical AV-DD mutant (AV1806DD), this mutant is limited in its utility as it results in constitutive unfolding of the R9 domain, which would also prevent talin regulation by autoinhibition. While this pursuit will be greatly facilitated by higher resolution analysis of the unique structural aspects of the talin-RIIα complex, more immediate empirical efforts may also be informed by analysis of talin mutants that are analogous to single nucleotide polymorphisms in known AKAPs that spare α-helix formation but significantly reduce PKA-binding (65). While there is clearly much work to be done, this report establishes an important new protein-protein interaction that impacts our understanding of PKA localization, talin function, and mechanotransduction.

## Materials and Methods

### Plasmids and cloning

To generate RIIα-mTb cell lines, a high efficiency and low background recombination-mediated cassette exchange (HILO-RMCE) method (94) was used, which utilize the Cre-Lox system. The HILO-RMCE donor plasmid (pEM791) and pCAGGS-nlCre were gifts from Dr. E. V. Makeyev (94). Plasmids expressing murine PKA (mPKA) RIIα and ER membrane targeting signal fused miniTurbo promiscuous biotin ligase (ER-mTb) were obtained from Addgene (#45527 and #107174, respectively). The mPKA RIIα sequence was PCR amplified and cloned into the upstream of the mTb sequence, replacing the ER membrane targeting signal and having a 26 amino acid linker between RIIα and mTb (pCDNA3-mPKA RIIα-mTb-V5), and then the mPKA RIIα-mTb sequence was cloned into the downstream of tetracycline responsive element (TRE) on pEM791 plasmid (pEM791-mPKA RIIα-mTb-V5) using Gibson assembly. The I3S, I5S mutant (mt) RIIα-mTb-V5 plasmid was generated by site directed mutagenesis (QuikChange II XL, Agilent) based on the wild type (WT) plasmid.

The plasmid expressing the recombinant PKA RIIα (ABK42343.1, full length) based on pET28b vector was obtained from Dr. J.D. Scott (University of Washington). A DNA fragment, which includes V5 tag-C3 protease site-His tag sequences, was synthesized (TWIST Bioscience) and cloned into the downstream of the RIIα sequence using Gibson assembly (pET28b-PKA RIIα FL-V5-C3-His).

For recombinant talin1 fragments, the cDNAs encoding murine talin1 residues, 482-911 (R1-R3), 913-1653 (R4-R8), 1655-2294 (R9-12) and 2300-2541(R13-DD) were synthesized by PCR using a mouse talin 1 cDNA as template and cloned into the expression vector pET151 TOPO (Invitrogen), which includes His-tag and V5 epitope at the upstream of the TOPO cloning site. Then, to be used for RII overlay, V5 tag sequences were removed by self-ligation of PCR amplicons, which copied whole plasmid sequences, except for the V5 tag sequence. Plasmids for the truncated and helix50 mutant talin1 fragments used in RII overlay; R9-11 (1655-2140), R9-10 (1655-1975), R9 (1655-1826) and R9-12 (V2087P, V2087S and V2087A) were generated by directed mutagenesis (Agilent) using the R9-12 plasmid as the template. The truncated fragments were made by replacing the residues, K2141, Q1976 and M1827 with a stop codon, respectively. R9-12ΔR10 (1655-1820, 1970-2294), R10 (1821-1975) and helix41 mutants (V1807A, V1800A/V1807A and A1806D/V1807D) were generated either assembling PCR amplicons copied the corresponding sequences from the R9-12 plasmid or replacing WT sequences with DNA fragments (TWIST Bioscience) that include the corresponding mutations, respectively, using Gibson assembly.

### RIIα-mTb cell lines and transfection

The U2OS HILO-RMCE acceptor cell line was gifted from Dr. E.V. Makeyev (94). The acceptor cells were routinely propagated and maintained in DMEM/high-glucose medium supplemented with 10% FBS in presence of 3 μg/ml blasticidin S. To generate RIIα-mTb cell lines, expressing either WT or mt (I3S, I5S) mPKA RIIα fused mTb-V5 proteins, in a tetracycline dependent manner, the acceptor cells were co-transfected with either pEM791-PKA RIIα-mTb-V5 or pEM791-PKA RIIα (I3S, I5S)-mTb-V5 and pCAGGS-nlCre. To transfect one well of a 6-well plate, 1.5 μg of total DNA (1:10 ratio of the Cre plasmid to the fusion protein plasmid) were mixed with 6 μl of Fugene 6 transfection reagent (Promega) in 100 μl of Opti-MEM, following manufacturer’s protocol. Cells were incubated with the transfection mixture overnight, and then the medium was changed. At 48-hour post-transfection, transfected cells were selected by a three-step selection protocol, beginning with 2.5 μg/ml of puromycin for 48 h, followed by 5.0 μg/ml treatment for 4 days until the puromycin-sensitive cells were eliminated, and then selected cells were expanded and maintained in the presence of 2.5 μg/ml puromycin. For cells transiently expressing RIIα-mTb, ERm-mTb, or EGFP-talin1, U2OS cells were transfected with 1.5 μg of corresponding pCDNA3 or pEGFP based plasmids using Fugene 6 reagent per one well of a 6-well plate and harvested at the 48-hour post-transfection for the immunoblotting analysis.

### Focal Adhesion Cytoskeletal (FACS) complex preparation

FACS complexes were prepared using a modified method from Dr. M. Humphries (95). Briefly, cells were plated on 20 μg/ml Human plasma fibronectin (Millipore) coated plates. After ∼21 h, when reaching ∼70-80% confluency, cells were fixed with 6 mM of the reversible crosslinker, Dimethyl dithiobispropionimidate (DTBP) (ThermoFisher) at 37°C for 15 min, followed by washing with Ca^2+^ and Mg^2+^ containing phosphate-buffered saline (PBS). The fixed cells were briefly soaked in ice-cold modified RIPA buffer for 2 min (50 mM Tris-HCl pH 7.5, 150 mM NaCl, 5 mM EDTA, 1% IGEPAL CA-630, 0.5% Na deoxycholate) with protease inhibitor cocktail (Halt, ThermoFisher) and phosphatase inhibitors (10 mM NaF and 1 mM Na_3_VO_4_). Cell bodies and organelles were then washed out by 150 ml/s water flow, and the left FACS complexes were collected by scrapping in the adhesion recovery solution (125 mM Tris-HCl pH 6.8, 1% SDS and 150 mM DTT) and reverse-crosslinked at 37°C for 30 min. The FACS proteins were acetone precipitated, resuspended in a reduced DTT (25 mM) adhesion recovery buffer and immediately used for FA AKAP screening or stored at −20°C for further immunoblot or RII overlay analyses.

### Focal adhesion AKAP screening

RIIα-mTb cells were raised in the absence or presence of 2 μg/ml doxycycline for 18 h until the cell density reached ∼70% confluency, followed by incubation with 50 μM biotin-containing media for 3 h. FACS proteins were collected from the non-or Dox-treated cells and diluted in RIPA buffer (Millipore). Biotinylated proteins were isolated by streptavidin pull-down (SA-Pd) using SA-coated magnetic beads (Invitrogen). The biotin-labeled proteins were separated by SDS-PAGE, trypsinized, and analyzed by LC/MS-MS (UVM proteomics core).

### Antibodies and immunoblotting

Antibodies against talin and vinculin were obtained from Sigma Aldrich (T3287 and V9131, respectively). V5 antibody and Mouse IgG isotype control were from Invitrogen (R960-25 and 02-6502, respectively). Antibodies against PKA RIIα (Sc-909, Santa Cruz Biotechnology), AKAP79 (610314, BD Biosciences), peroxiredoxin-3 (LF-PA0255, Abfrontier), phospho-PKA substrates (9624, Cell Signaling Technology) were used in this study. HRP-conjugated streptavidin (SA-HRP) was purchased from ThermoFisher. For immunoblotting analysis, whole-cell extracts (WCE) were prepared by lysing cells in ice-cold RIPA lysis buffer (Millipore). WCE, FACS preparations, immunoprecipitations or SA-Pd sample proteins were separated by SDS-PAGE, transferred to PVDF membranes, blocked with either 5% BSA or 5% NFDM, and then immunoblotted with indicated antibodies or SA-HRP.

### Immunoprecipitation and pull-down assays

For immunoprecipitation (IP), RIIα-mTb cells were grown in the absence or presence of 2 μg/ml Dox for 18 h and then either non-(V5 co-IP) or treated with 50 μM biotin for 3 h (talin IP). For V5 IP, cells were further crosslinked with 6 mM DTBP at 37°C for 15 min. Cells were lysed in ice-cold modified RIPA buffer (20 mM Tris-HCl pH 7.5, 100 mM NaCl, 1 mM EDTA, 1% IGEPAL CA-630, 10% glycerol). 5 μM cytochalasin D and 2.5 μM Latrunculin B were added to cell lysates to disrupt polymerized actin, and the cell lysates were mixed with mouse IgG, talin or V5 antibodies and rocked end-over-end at 4°C overnight, followed by incubation with protein A/G-agarose beads (Santa Cruz Biotechnology) at 4°C for 4 h. Beads were then washed 4 times for 2 min with modified RIPA buffer. For streptavidin pull-down (SA-Pd), WT or mutant (I3S, I5S) RIIα-mTb-expressing cells were grown in the presence of 2 μg/ml Dox and incubated with 50 μM biotin containing media for 3 h. For blebbistatin treatment, the drug was added during this 3-hour biotin treatment at a final concentration, 25 μM. Cells were lysed as described for the IP samples and the lysate was rocked end-over-end with 50 μl of SA-coated magnetic beads (Invitrogen) at 4°C for 2 h. The beads were then washed thrice with cold, modified RIPA. For GFP trap, EGFP-talin1 transfected U2OS cells were re-seeded from 10 cm dishes to 15 cm, fibronectin (20 μg/ml) -coated plates at 48 h post transfection and grown for 24 h. Next, cells were incubated in either DMSO or forskolin/IBMX (25 μM/ 50 μM) containing media for 10 min and lysed in RIPA buffer (Cold Spring Harbor). The cell lysate was diluted in 10 mM Tris-HCl, pH 7.5, 150 mM NaCl, 0.5 mM EDTA and end-to-end rotated with 20 μl of GFP trap magnetic beads (ChromoTek) at 4°C for 1 h. The beads were then washed four times with cold dilution buffer with 0.05% IGEPAL CA-630. For IP, SA-Pd and GFP trap assays, all RIPA and dilution buffers were supplemented with protease inhibitor cocktail (Halt, ThermoFisher) and phosphatase inhibitors (10 mM NaF and 1 mM Na_3_VO_4_), and bead-bound proteins were eluted by boiling in 2X Laemmli sample buffer, then stored at −20°C or analyzed immediately by immunoblotting.

### Protein expression and purification

For overlay assays, his-tagged talin1 construct plasmids and pET28b-PKA RIIα FL-V5-C3-His were transformed into *Escherichia coli* BL21(DE3)*, and then protein expression was induced with 1 mM IPTG at 37°C for 3 h. For purification, cells containing the induced proteins were either lysed by lysozyme-containing buffer (50 mM sodium phosphate, pH 7.5, 100 mM NaCl, 2 mM MgCl_2_, 0.5% triton-X, 4 mg/ml lysozyme and Halt protease cocktail) or sonicated in Tris buffer (10 mM Tris-HCl, pH 7.5, 150 mM NaCl and Halt protease cocktail). His-tagged recombinant proteins were isolated by 1 h incubation with Talon metal affinity resin (Clontech) at room temperature. Beads were washed 3 times either with 50 mM sodium phosphate or 10 mM Tris-HCl, containing 500 mM NaCl and 0.5% Triton-X 100, pH 7.5, followed by 3 times second wash with buffer with no detergent and low salt (100 mM NaCl), and then the recombinant proteins were eluted with the second wash buffer containing 150 mM imidazole.

For PKA RIIα FL-V5 proteins, His6 tags were cleaved by C3 protease (GenScript), according to the manufacturer’s protocol. Briefly, the talon bead eluate was buffer-exchanged to 10 mM Tris-HCl, pH 7.5, 50 mM NaCl using 10k molecular weight cut-off column (Millipore) to remove imidazole and incubated with C3 protease for 16 h at 4°C. The C3 proteases, which are His-tagged, were removed by the talon resin, and the supernatant that includes the His-tag cleaved RIIα FL-V5 proteins was collected and concentrated.

For NMR assays, pET151 plasmids for mouse talin1 R9 (residues 1655-1822), R10 (residues 1815-1973), R9-R10 (residues 1655-1973), were used as reported previously (64). Additional codon optimized synthetic genes encoding, h41-R10 (residues 1785-1973), talin2 h41-R10 (1792-1974), and talin1 R9-h42 (1655-1841) were produced by GeneArt. The truncated helix42 constructs, Talin1 R9-h42L (1785-1835) and Talin1 R9-h42S (1785-1828) plasmids were made by site-directed mutagenesis using Talin1 R9-h42 plasmid. To produce ^15^N-labeled proteins, talin constructs were transformed into BL21(DE3)* E. coli cells and grown in 10 mL of LB + 100 mg/L ampicillin at 37°C overnight as starter cultures and then in 2M9 minimal medium which contains ^15^N-ammonium chloride and 100 mg/L ampicillin at 37°C till OD_600_ of 0.7-0.8. After inducing cells with 1 mM IPTG, the cultures were incubated overnight at 20°C. The following day, harvested cells pellets were resuspended in 20 mM Tris-HCl, pH 8, 500 mM NaCl, and 20 mM imidazole, and lysed by sonication. The proteins were purified by nickel affinity chromatography using a 5 ml HisTrap HP column (GE Healthcare). The eluted proteins were dialyzed into 20 mM Tris-HCl, pH 8, 50 mM NaCl with AcTEV protease (Invitrogen) to remove the His-tag. After overnight dialysis, the proteins were purified with a HiTrap Q HP cation exchange column (GE Healthcare). Further details are available elsewhere (96).To produce unlabeled proteins, cells were grown in LB + appropriate antibiotics (100 μg/ml ampicillin for talin constructs and 50 μg/ml kanamycin for PKA RII-DD) at 37°C. Once OD_600_ 0.7-0.8, cells were induced and harvested, and the proteins were purified in the same way as ^15^N-labeled proteins except no TEV cleavage for PKA RII-DD protein.

### RII overlay

1.5 μg of recombinant proteins or 10 μg of the U2OS WCE or FACS proteins were separated by SDS-PAGE and transferred to nitrocellulose membranes. Membranes were blocked with 10% non-fat dry milk (NFDM) in TBS-T for 1 h and then incubated with either PKA RII-DD-V5 or full-length PKA RIIα-V5 solution (0.2 μg/ml concentration in 10% NFDM, containing either 0.4 μM Ht31p or Ht31 peptides) for 12 h. After extensive washing with TBS-T, membranes were incubated with V5-HRP-containing solution (BioLegend, 1:4000, 10% NFDM) for another 12 h, washed with TBS-T, and developed by enhanced chemiluminescence.

### NMR analysis

^15^N-labeled protein samples were prepared at 150 μM final concentration in 12 mM NaH_2_PO_4_, 6 mM Na_2_HPO_4_, pH 6.5, 50 mM NaCl, 2 mM DTT, 5%(v/v) D_2_O. NMR spectra were collected at 298 K on Bruker Avance III 600 MHz NMR spectrometer equipped with CryoProbe. All data was processed via TopSpin and analyzed with CCPN Analysis (97).

### Immobilized protein extension assay

A modification of a published immobilized protein extension (IPE) assay (71) was used. Briefly, a flexible silicon sheet (PDMS KRN Silicone Film (KRN-300um-10×12cm), AliExpress) was held taught in a 4” crochet hoop (Item # 10124139, Michaels), fiduciary marks were made with a lab marker (to visualize the degree of membrane stretch; see below), and the sheet was exposed to room air plasma (PDC-32G, Harrick Plasma) for 30 sec. The sheet was immediately coated with 0.5 ml of 10 μg/ml purified talin1 R9-12 fragment to allow passive adsorption to the PDMS surface. After washing and blocking with BSA, the talin1-coated sheet was left unstretched or lowered to varying degrees over an inverted, 70 mm diameter flat-bottom crystallization dish (PYREX™ 314070, Fisher Scientific) to stretch the immobilized talin fragments then incubated with 0.2 μg/ml PKA RIIα-D/D. All coating and binding steps were performed at RT. Following extensive washing, proteins bound to the sheets were collected in hot Laemmli sample buffer and analyzed by immunoblotting.

### Single-molecule manipulation

The single-molecule manipulation experiments were carried out as described previously (68), using a custom high-force magnetic tweezers platform that can exert forces up to 100 pN with ∼1 nm extension resolution for immobilized beads at a 200 Hz sampling rate (69, 70). For the unfolding experiments, the protein of interest was immobilized on the glass coverslip of a laminar flow chamber and tethered to a 3-mm paramagnetic bead (Dynabeads M270 streptavidin) using Halo-tag/Halo-ligand and biotin/streptavidin chemistry. Briefly, glass coverslip was cleaned in an ultrasonic cleaner in 10% Decon 90 solution, followed by acetone and isopropanol for 30 min each. Then the coverslips was silanized by 1% APTES (Sigma-Aldrich) in methanol for 20 min and rinsed clean by methanol. The APTES-coated coverslip was assembled into a flow channel and NH2-O4-Halotag ligand (Promega) was immobilized on to the coverslip through glutaraldehyde (Sigma-Aldrich) crosslinking. The channel was blocked by 1M Tris-HCl, pH 7.4 for 30 min followed by 1% BSA in 1X PBS and 0.1% Tween-20 overnight. The stretchable talin1 R9-12 protein containing Halo-tag and biotinylated Avi-tag (described previously (68)) was immobilized by flowing ∼0.1 mg/ml protein into the channel for 20 min. And then streptavidin-coated M270 beads were added to the channel to form the tether. To avoid flow-induced unfolding of the immobilized tether during solution exchange, a buffer isolation membrane well array (98) was implemented. All unfolding and refolding experiments were carried out in 1X PBS, 1% BSA, 1 mM TCEP and 0.1% Tween-20.

## Acknowledgements

We thank Eugene Makeyev (King’s College London) and John Scott (University of Washington) for cell lines and plasmids. The authors are indebted to Anna Schmoker (Director, Targeted Protein Degradation Proteomics Core, Dana-Farber Cancer Institute) for phosphoproteomic analyses of talin1 fragments. The work reported here was supported by funds from the University of Vermont Cancer Center and NIH grant R01GM137611 to A.K.H. and BBSRC grant BB/S007245/1 and Cancer Research UK Program grant DRCRPG-May21 to B.T.G, National Research Foundation (NRF), Prime Minister’s Office, Singapore under its NRF Investigatorship Programme (NRF Investigatorship Award No. NRF-NRFI2016-03) and grants from the National Research Foundation through the Mechanobiology Institute Singapore to J.Y.

