## Supplemental Figures and Table for "The focal adhesion protein talin is a mechanically-gated A-kinase anchoring protein (AKAP)"

### Supporting Information Text

#### SI Methods and Materials

*In vitro kinase assay.*

The catalytic subunit of PKA (PKAc) was purchased from New England Biolabs (NEB), and the kinase reaction was performed according to the manufacture's protocol. Briefly, 12 µg of talin R4-8 or R9-12 fragments was mixed with 2500 units of PKAc in 1X protein kinase buffer (NEB) and incubated at 30 °C for 1 hours in the absence or presence of 200 µM ATP. Reaction products were analyzed by SDS-PAGE and western blotting with anti-phospho-PKA substrate antibody as described in the main text.

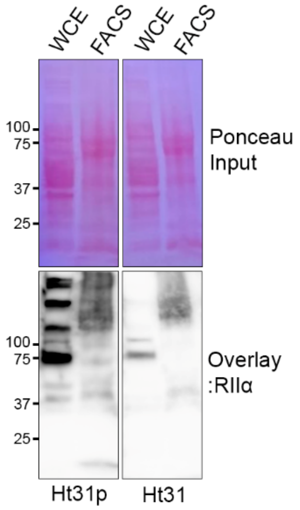


**Fig. S1. Screening for Focal Adhesion AKAPs.** PKA RIIα D/D domain overlay of whole-cell extract (WCE) and focal adhesion/cytoskeletal (FACS) proteins from U2OS cells in the presence of Ht31p and Ht31, inactive and competitive inhibitor peptides for RII D/D binding, respectively.

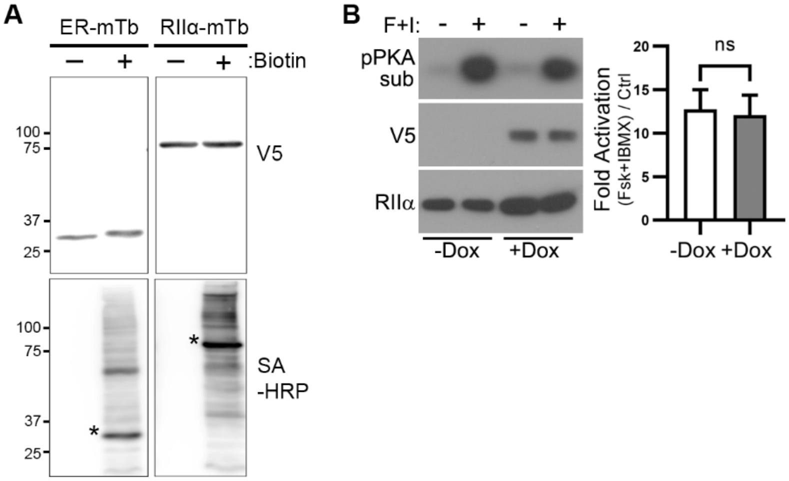


**Fig. S2. Characterization of PKA RIIα-miniTurbo biotin ligase fusion protein. (*A*)** A V5 epitope-tagged miniTurbo biotin ligase fused to either an ER membrane targeting sequence (*ERm-mTb*) or to PKA RIIα (*RIIα-mTb*) were transiently expressed in U2OS cells. Cells were labelled with biotin for 3 h and WCE were analyzed by blotting for fusion protein expression using V5 antibody and protein biotinylation using streptavidin-HRP (*SA-HRP*). Asterisks denote ligase self-biotinylation. **(*B*)** Expression of RIIα-mTb does not alter PKA activity. (*Left*) U2OS cells stably expressing V5-tagged RIIα-mTb under an inducible promoter were treated (or not) with doxycycline (*Dox*) to induce expression then treated with forskolin & IBMX to activate PKA. Lysates were analyzed by blotting for RIIα or V5, or by *in vitro* PKA assay using purified GFP-R4S protein as a substrate. Reaction products were analyzed by blotting with a phospho-PKA substrate antibody. (*Right*) Average fold-activation (-/+s.d.) for n=3 experiments is shown.

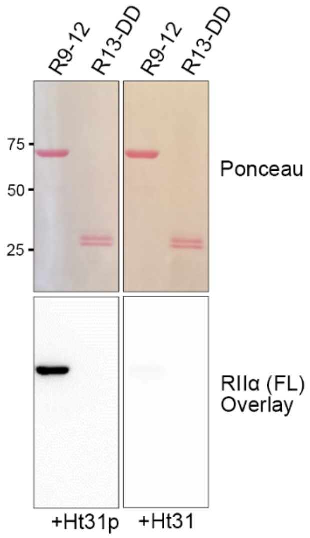


**Fig. S3. Direct binding of full-length PKA RIIα to talin1 R9-12.** Recombinant fragments comprising talin1 R9-12 or R13-DD (dimerization domain) were separated by SDS-PAGE and transferred to membranes which were stained to ensure equal loading (*Ponceau*) then analyzed for direct PKA interaction by overlay with purified, full-length RIIα subunits (*RIIα FL*) in the presence of a competitive peptide that blocks AKAP binding (*Ht31*) or a non-blocking negative control peptide (*Ht31p*).

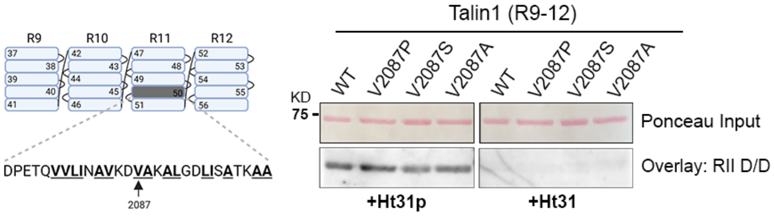


**Fig. S4. The *in silico* predicted PKA RII binding site, Helix-50, is not required for the RII-Talin interaction.** (*Left*) A schematic depicting talin fragment containing R9-12, highlighting the highest-scoring *in-silico* predicted AKAP consensus sequence in helix50 within R11 (hydrophobic residues of the amphipathic helix, predicted to bind PKA R subunits, are underlined; Val2087, numbered as in full-length talin, is indicated for reference). (*Right*) Three talin R9-12 fragments with point mutations targeting Val2087 in the middle of the predicted AKAP consensus sequence, were analyzed, along with the wild-type (WT) R9-12 fragment, by overlay assay with the PKA RII D/D domain in the presence of Ht31p or Ht31 control or competitive inhibitor peptide.

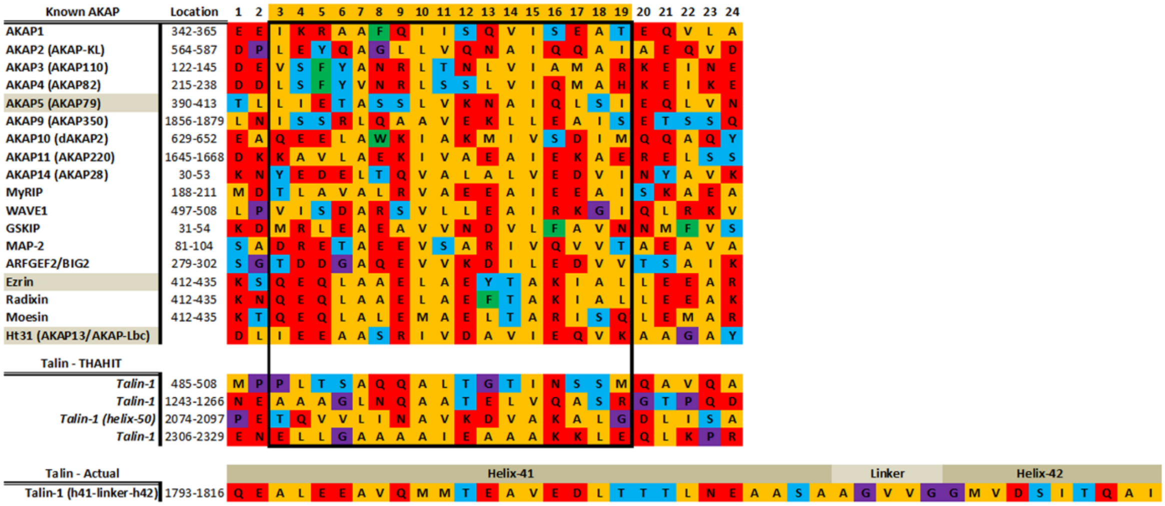


**Fig. S5. Sequence alignment of the PKA RIIα-binding motif in known AKAPs and comparison to predicted and experimentally determined motifs in talin1.**

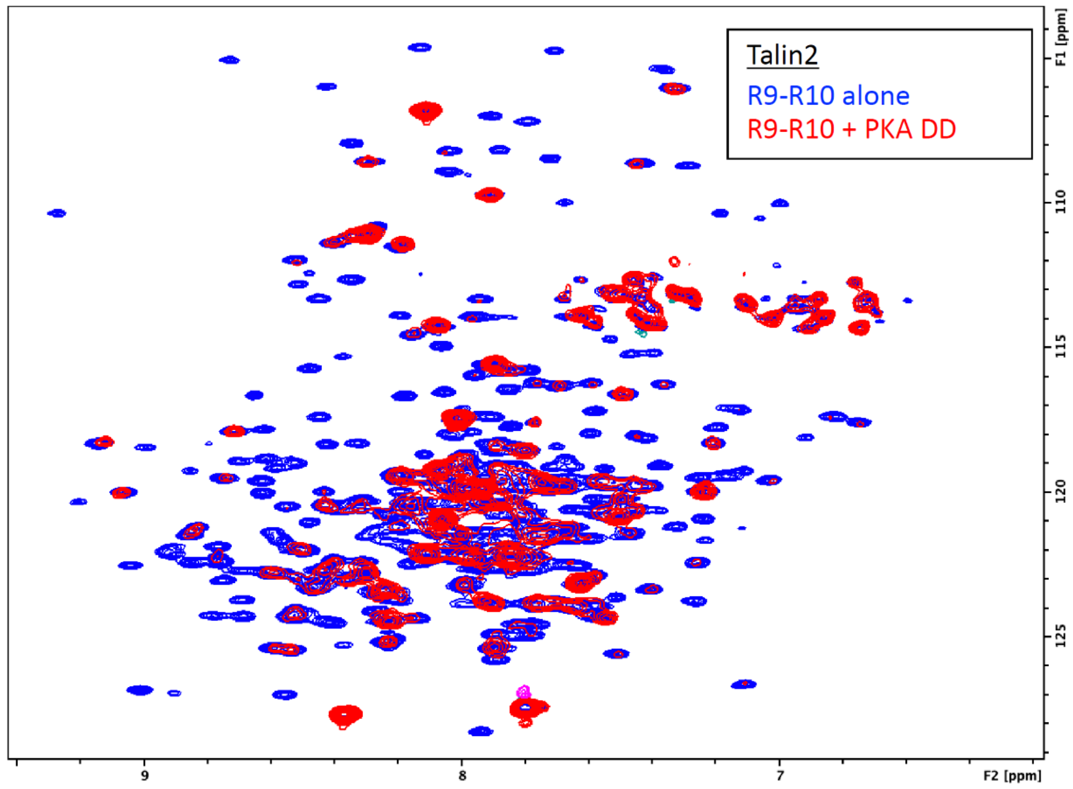


**Fig. S6. PKA RIIα D/D binds Talin2 R9-R10.** HSQC-spectra of talin2 R9-R10 (*blue*) and upon addition of the PKA RIIα D/D domain (*red*). As seen with talin1, the peaks that broaden out upon addition of the RIIα D/D domain are almost exclusively from R9.

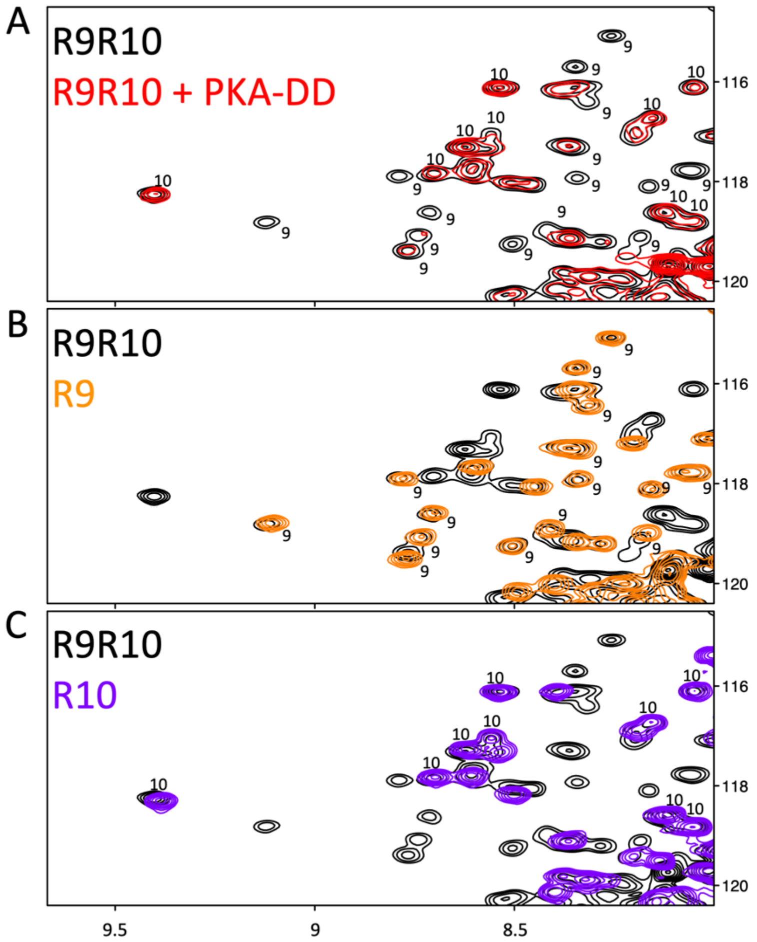


**Fig. S7 Talin R9 domain unfolds in the R9-R10 fragment upon PKA RIIα-D/D addition. (*A*)** HSQC-spectra of talin1 R9-R10 alone (*black*) and upon addition of PKA RIIα-D/D domain (*red*). The peaks are assigned "9" or "10" depending on which domain they pertain to. The peaks that are broadened out are almost exclusively from R9. **(*B-C*)** Overlay of R9 (*B**,** orange*) and R10 (*C**,** blue*) with R9-R10 (*B* & *C*, *black*) to show how the peaks in (*A*) were unambiguously assigned.

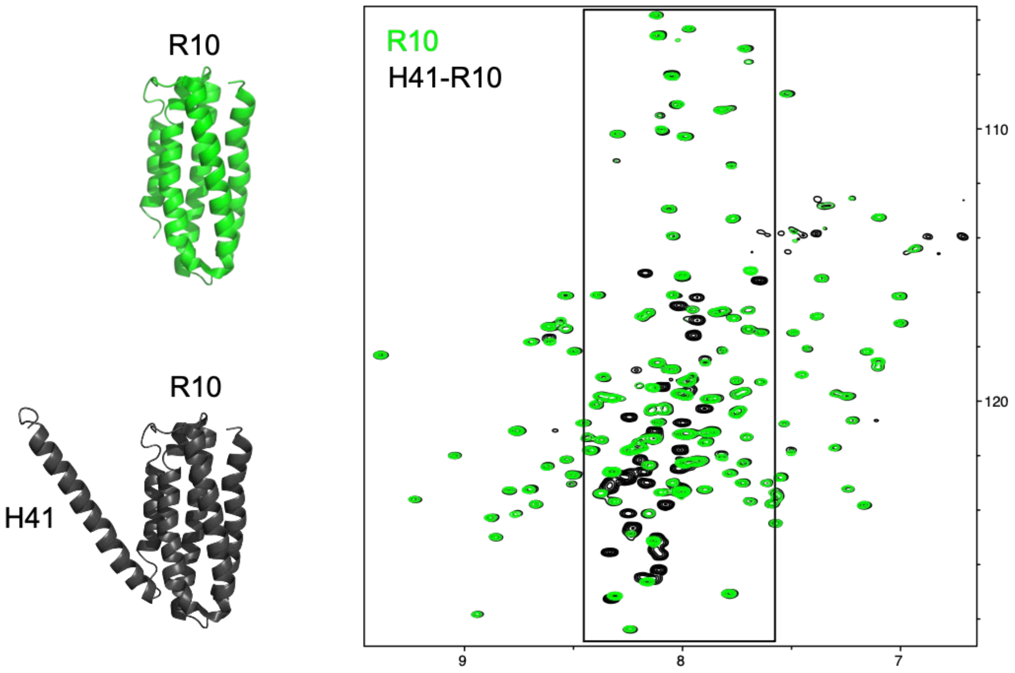


**Fig. S8 Comparison of NMR spectra of talin R10 and h41-R10 allows identification of h41-specific peaks.** $^{1}H$,$^{15}N$-HSQC spectra of $^{15}N$-labeled talin1 R10 (residues 1815-1973; *green*) and helix41-R10 (h41-R10; residues 1792-1973; black), the signals from h41 are clearly visible as unfolded peaks in the central boxed region of the spectra.

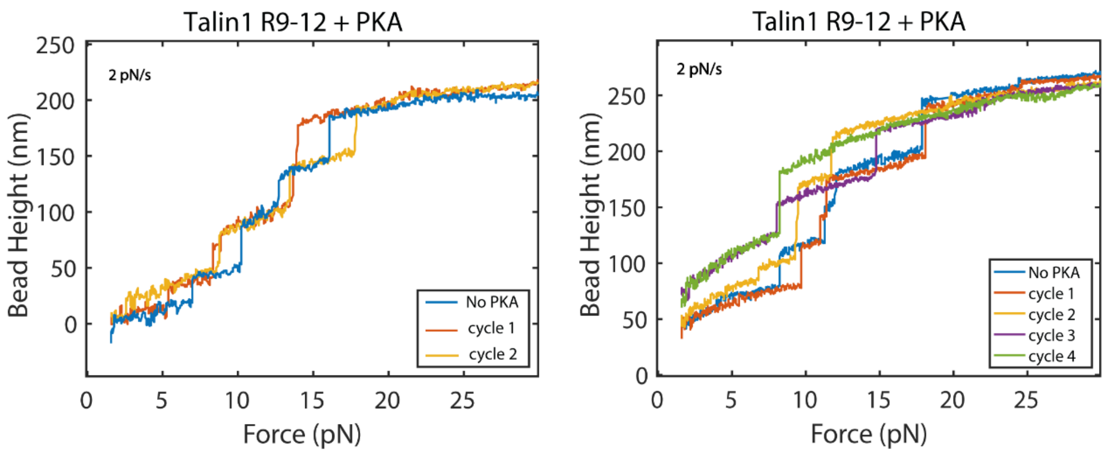


**Fig. S9. Force-extension cycles of talin1 R9-12 in the absence and presence of PKA RIIα D/D domain.** Two independent experiments showing force-extension curves of paramagnetic bead-tethered talin1 R9-12 before and after addition of 100 nM PKA RIIα D/D domain.

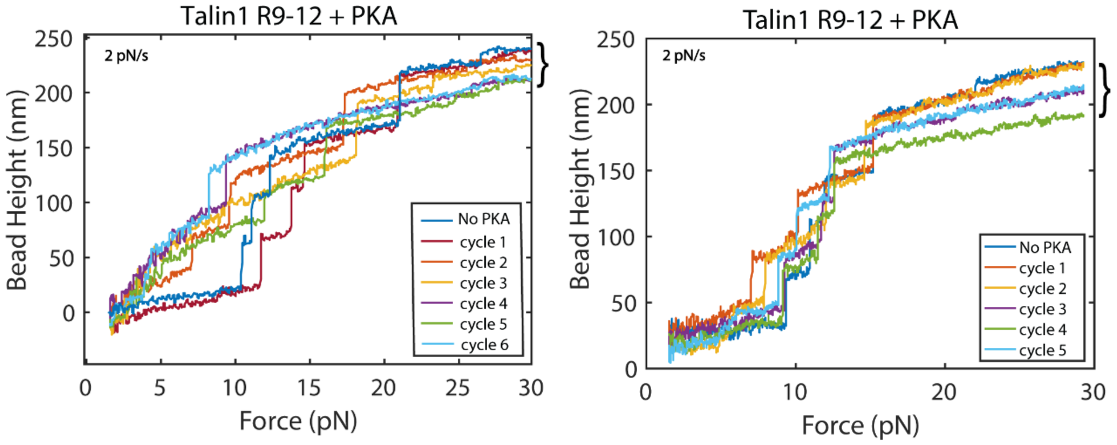


**Fig. S10. Force-extension cycles of talin1 R9-12 in the absence and presence of PKA RIIα D/D domain.** Two independent experiments showing force-extension curves of paramagnetic bead-tethered talin1 R9-12 before and after addition of 100 nM PKA RIIα D/D domain, highlighting the reduction in the maximum extension length of R9-12 at maximal force (*brackets*) after increased numbers of force loading cycles in the presence of RIIα D/D. Note that the graph on the left represents the same experiment as shown in Fig. 5B, but depicts a larger number of cycles.

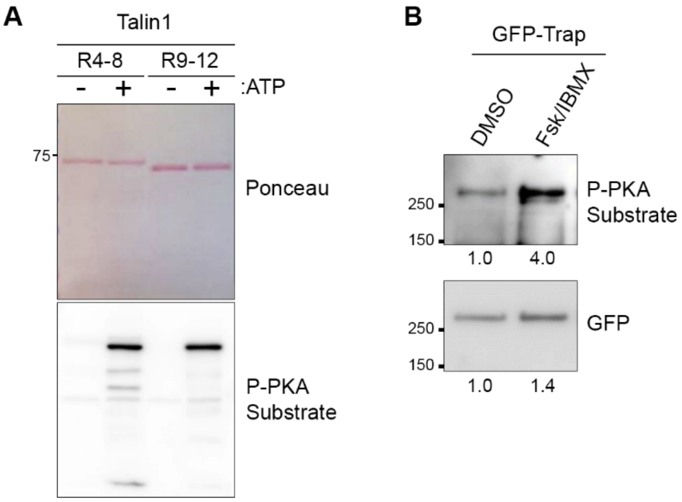


**Fig. S11. Phosphorylation of talin1 by PKA. (*A*)** Talin1 R4-8 or R9-12 fragments were incubated with PKA catalytic subunits in the absence or presence of ATP *in vitro*, then the phosphorylation of talin fragments was tested by immunoblotting with a phospho-PKA substrate antibody. **(*B*)** GFP-tagged talin1 was transfected into U2OS cells which were treated with solvent (DMSO) or forskolin and IBMX (Fsk/IBMX, 25 μM/50 μM) for 10 min. Transfected talin1 was isolated by GFP-trap and phosphorylation was measured by immunoblotting with phospho-PKA substrate antibody.

**Table S1.** Summary of PKA binding of talin fragments.

| Construct | Sequence (*Reference* / *PKA Binding* / *Minimal Binding* / *No PKA Binding*) |
|---|---|
| **h41-*LINKER*-h42** | 1785 NPKQAAHTQEALEEAVQMMTEAVEDLTTTLNEAASA*AGVVGG*MVDSITQAINQLDEG 1841 |
| **R9-R10** | **R9**-NPKQAAHTQEALEEAVQMMTEAVEDLTTTLNEAASAAGVVGGMVDSITQAINQLDEG-**R10** |
| **R9** | **R9**-NPKQAAHTQEALEEAVQMMTEAVEDLTTTLNEAASAAG |
| **R10** | NEAASAAGVVGGMVDSITQAINQLDEG-R10 |
| **h41-R10** | NPKQAAHTQEALEEAVQMMTEAVEDLTTTLNEAASAAGVVGGMVDSITQAINQLDEG-**R10** |
| **R9-h42** | **R9**-NPKQAAHTQEALEEAVQMMTEAVEDLTTTLNEAASAAGVVGGMVDSITQAINQLDEG |
| **R9-h42-long** | **R9**-NPKQAAHTQEALEEAVQMMTEAVEDLTTTLNEAASAAGVVGGMVDSITQAI |
| **R9-h42-short** | **R9**-NPKQAAHTQEALEEAVQMMTEAVEDLTTTLNEAASAAGVVGGMV |
| **AV1806DD** | **R9**-NPKQAAHTQEALEEAVQMMTE**DD**EDLTTTLNEAASAAGVVGGMVDSITQAINQLDEG-**R10** |
